# Mutations of SARS-CoV-2 nsp14 exhibit strong association with increased genome-wide mutation load

**DOI:** 10.1101/2020.08.12.248732

**Authors:** Doğa Eskier, Aslı Suner, Yavuz Oktay, Gökhan Karakülah

## Abstract

SARS-CoV-2 is a betacoronavirus responsible for human cases of COVID-19, a pandemic with global impact that first emerged in late 2019. Since then, the viral genome has shown considerable variance as the disease spread across the world, in part due to the zoonotic origins of the virus and the human host adaptation process. As a virus with an RNA genome that codes for its own genomic replication proteins, mutations in these proteins can significantly impact the variance rate of the genome, affecting both the survival and infection rate of the virus, and attempts at combating the disease. In this study, we analyzed the mutation densities of viral isolates carrying frequently observed mutations for four proteins in the RNA synthesis complex over time in comparison to wildtype isolates. Our observations suggest mutations in nsp14, an error-correcting exonuclease protein, have the strongest association with increased mutation load in both regions without selective pressure and across the genome, compared to nsp7, 8, and 12, which form the core polymerase complex. We propose nsp14 as a priority research target for understanding genomic variance rate in SARS-CoV-2 isolates, and nsp14 mutations as potential predictors for high mutability strains.

## Introduction

COVID-19 is an ongoing global pandemic characterized by long-term respiratory system damage in patients, and caused by the SARS-CoV-2 betacoronavirus. It is likely of zoonotic origin, but capable of human-to-human transmission, and since the first observed cases in the Wuhan province of China (Chan et al., 2020; Riou & Althaus, 2020), it has infected over 14 million people, with 612054 recorded deaths. In addition to its immediate effects on the respiratory system, its long term effects are still being researched, including symptoms such as neuroinvasion (Li, Bai & Hashikawa, 2020; Wu et al., 2020), cardiovascular complications (Kochi et al., 2020; Zhu et al., 2020), and gastrointestinal and liver damage (Lee, Huo & Huang, 2020; Xu et al., 2020). Due to its high transmissibility, and capacity for asymptomatic transmission (Wong et al., 2020), study of COVID-19 and its underlying pathogen remain a high priority. As a result, the high amount of frequently updated data on viral genomes on databases such as GISAID (Elbe & Buckland-Merrett, 2017) and NextStrain (Hadfield et al., 2018) provides researchers with invaluable resources to track the evolution of the virus as it spreads across the world.

SARS-CoV-2 has a linear, single-stranded RNA genome, and does not depend on host proteins for genomic replication, instead using an RNA synthesis complex formed from nonstructural proteins (nsp) coded by its own genome. Four of the key proteins involved in the complex are nsp7, nsp8, nsp12, and nsp14, all of which are formed from cleavage of the polyprotein Orf1ab into mature peptides. Nsp12, also known as RdRp (RNA-dependent RNA polymerase), is responsible for synthesizing new strands of RNA using the viral genome as a template. Nsp7 and nsp8 act as essential co-factors for the polymerase unit, together creating the core polymerase complex (Kirchdoerfer & Ward, 2019; Peng et al., 2020), while nsp14 is an exonuclease which provides error-correcting capability to the RNA synthesis complex (Subissi et al., 2014; Ma et al., 2015; Romano et al., 2020). Owing to their role in maintaining replication fidelity and genome sequences, these proteins are key targets of study in understanding the mutation accumulation and adaptive evolution of the virus (Peng et al., 2020).

In our previous study, we examined the top 10 most frequent mutations in the SARS-CoV-2 nsp12, and identified that four of them are associated with an increase in mutation density in two genes, the membrane glycoprotein (M) and the envelope glycoprotein (E) (the combination of which is hereafter referred to as MoE, as we previously described), which are not under selective pressure, and mutations in these genes are potential markers of reduced replication fidelity (Eskier et al., 2020). In this study, we follow up on our previous findings and analyze the mutations in nsps 7, 8, and 14, in addition to nsp12, to identify whether the mutations are associated with a nonselective increase in mutation load or not. We then examine whole genome mutation densities in mutant isolates in comparison to wildtype isolates using linear regression models, in order to understand whether the mutations are associated with potential functional impact. Our findings indicate that mutations in nsp14 are most likely to be predictors of accelerated mutation load increase.

## Materials and Methods

### Genome sequence filtering, retrieval, and preprocessing

As previously described (Eskier et al., 2020), isolate genomes obtained from the GISAID EpiCoV database (date of accession: 17 June, 2020) were filtered to remove low-coverage or incomplete genomes, aligned against the reference genomic sequence for SARS-CoV-2, and processed to identify any SNVs present in the isolates and their impact on peptide sequences, if any, using the MAFFT, snp-sites, bcftools, and ANNOVAR suite of software. Following the alignment and variant annotation, isolates with incomplete sequencing location or date data were further removed from the pool, and the 5’ UTR (bases 1-265) and the 100 nucleotides at the 3’ end of the genome were masked due to their gap-heavy and low-quality nature. Following the filters, 29,600 genomes were used for the analyses.

### Mutation density calculation

Variants were categorized as synonymous and nonsynonymous following annotation by ANNOVAR, with intergenic or terminal mutations being considered synonymous. Gene mutation densities were calculated separately for synonymous and nonsynonymous mutations, as well as the total of SNVs, for each isolate, using a non-reference nucleotides per kilobase of region metric. Mutation densities were calculated for the combined membrane glycoprotein (M) and envelope glycoprotein (E) genes (MoE), the surface glycoprotein gene (S), and the whole genome.

### Statistical Analysis

Descriptive statistics for continuous variable days were calculated with mean, standard deviation, median, and interquartile range. Kolmogorov–Smirnov test was used to check the normality assumption of the continuous variables. In cases of non-normally distributed data, the Wilcoxon rank-sum (Mann-Whitney U) test was performed to determine whether the difference between the two MoE status groups was statistically significant. The Fisher’s exact test and the Pearson chi-square test were used for the analysis of categorical variables. The univariate logistic regression method was utilized to assess the mutations associated with MoE status in single variables, and then multiple logistic regression method was performed. The final multiple logistic regression model was executed with the backward stepwise method. The relationship between mutation density and time in isolates with mutations of interest, as well as in the group comprising all isolates, was examined via non-polynomial linear regression model and Spearman’s rank correlation. A p-value of less than 0.05 was considered statistically significant. All statistical analyses were performed using IBM SPSS version 25.0 (Chicago, IL, USA).

## Results and Discussion

### Increases in the mutation load of SARS-CoV-2 are unevenly distributed across its genome

To identify the trends in SARS-CoV-2 mutation load over time, we calculated the average mutation density per day for all isolates for whole genome, S gene, and MoE regions, capping outliers at the 95^th^ and 5^th^ percentile values to minimize the potential effects of sequencing errors (Fig. 1). Our results show that both at the genome level and the S gene, a very strong positive correlation between average mutation density and time. In comparison, MoE has a weak positive correlation, with a wider spread of mean density in early and late periods compared to the genome and the S gene. This is consistent with reduced selective pressure on the M and E genes, as has previously been described (Dilucca et al., 2020). The top nonsynonymous mutation is 23403A>G (in 22,271 isolates), responsible for the D614G substitution in the spike protein, followed by the 14408C>T mutation (in 22,226 isolates) in the nsp12 region of the Orf1ab gene, causing P323L substitution in the RdRp protein, and the 28144C>T mutation (in 3,081 isolates), responsible for the L84S substitution in the Orf8 protein. The most common synonymous mutation is the 8782C>T mutation (in 3,047 isolates), and is found on the nsp4 coding region of the Orf1ab gene. For the S gene, the most frequent synonymous mutation is the 23731C>T mutation (in 622 isolates), and the second most common nonsynonymous mutation, after the aforementioned D614G mutation, is 25350C>T (in 215 isolates), responsible for the P1263L substitution. For MoE, the most common synonymous and nonsynonymous mutations are 26735C>T (in 341 isolates) and 27046C>T (in 530 isolates), respectively, both of which are found in the M gene, and the latter of which causes T175M amino acid substitution. Other than the D614G mutation, all of the mentioned mutations are C>T substitutions, the prevalence of which in T- or A-rich regions of the SARS-CoV-2 genome have been previously documented (Simmonds, 2020).

**Figure 1.**
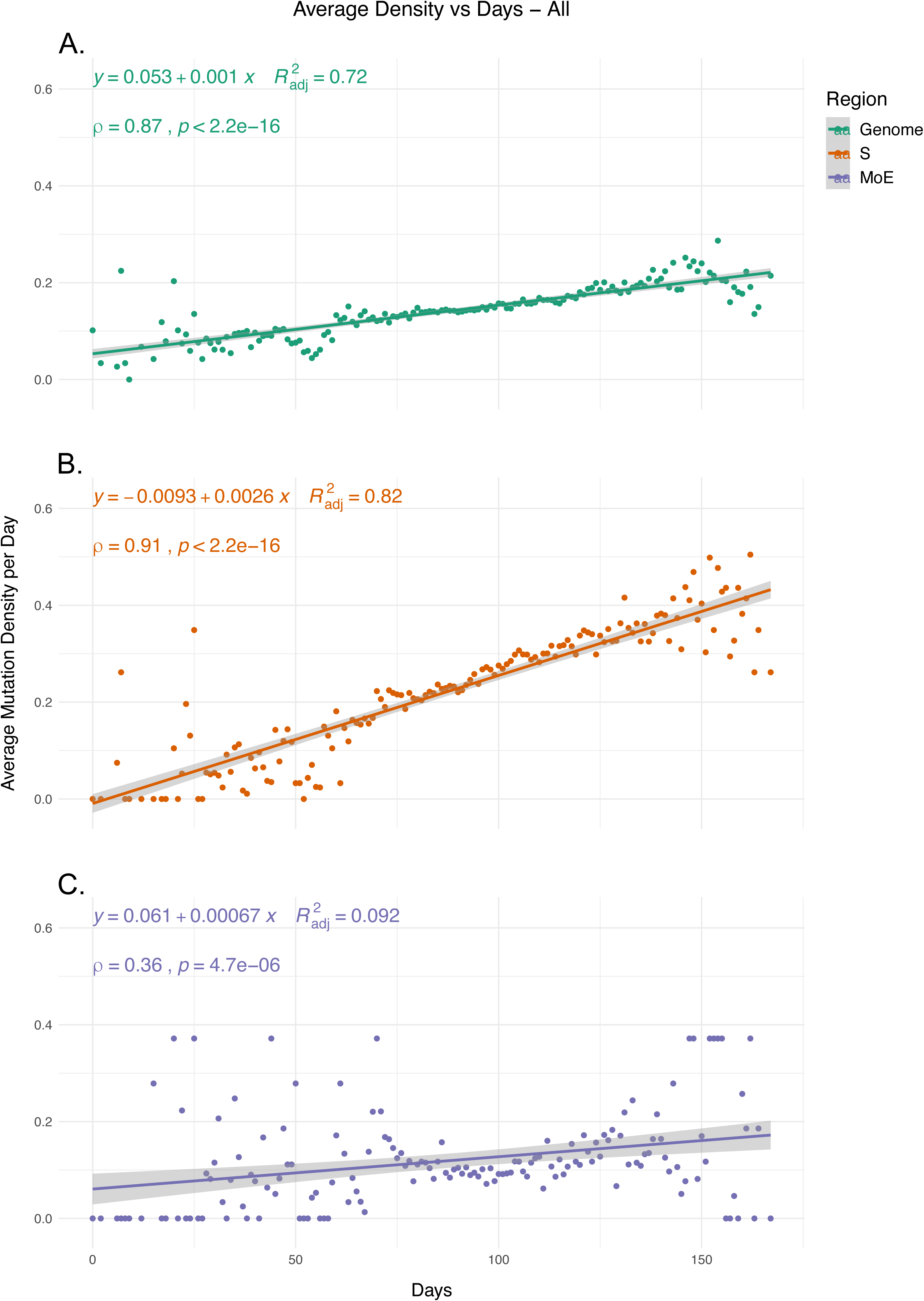
The average mutation density per day for genome, S gene, and M and E genes. Correlation scores are calculated using Spearman rank correlation.

### Mutations in RNA synthesis complex proteins are associated with higher mutation load

After identifying the increase in mutation load over time, which was more prominent in genes with high functional impact (S, Orf1ab) compared to other structural genes (M, E, N), we sought to examine possible associations of variants in proteins involved in SARS-CoV-2 genome replication with the increase. We first identified the five most frequently observed mutations for nsps 7, 8, 12 (also known as RdRp) and 14, four of the proteins cleaved from the Orf1ab polyprotein and are involved in the RNA polymerization, followed by analyzing the association of each mutation with the presence of MoE mutations (hereafter referred to as MoE status) using the chi-square test. 12 out of the 20 mutations were found to have a significant association with MoE status (p-value < 0.05) (Table 1). Compared to our previous findings on the top 10 nsp12 mutations (Eskier et al. 2020), which was based on an analysis of 11,208 samples as of 5 May 2020, 13536C>T and 13862C>T have increased in rank of appearance, from 6^th^ and 7^th^ to 4^th^ and 5^th^, respectively, and decreased in p-value to show statistically significant associations. In addition, the 13730C>T mutation have increased in rank of appearance from 4^th^ to 3^rd^. Out of the other nsps tested, nsp14 was found to have four significant mutations, while nsp7 had two and nsp8 had one.

**Table 1.**
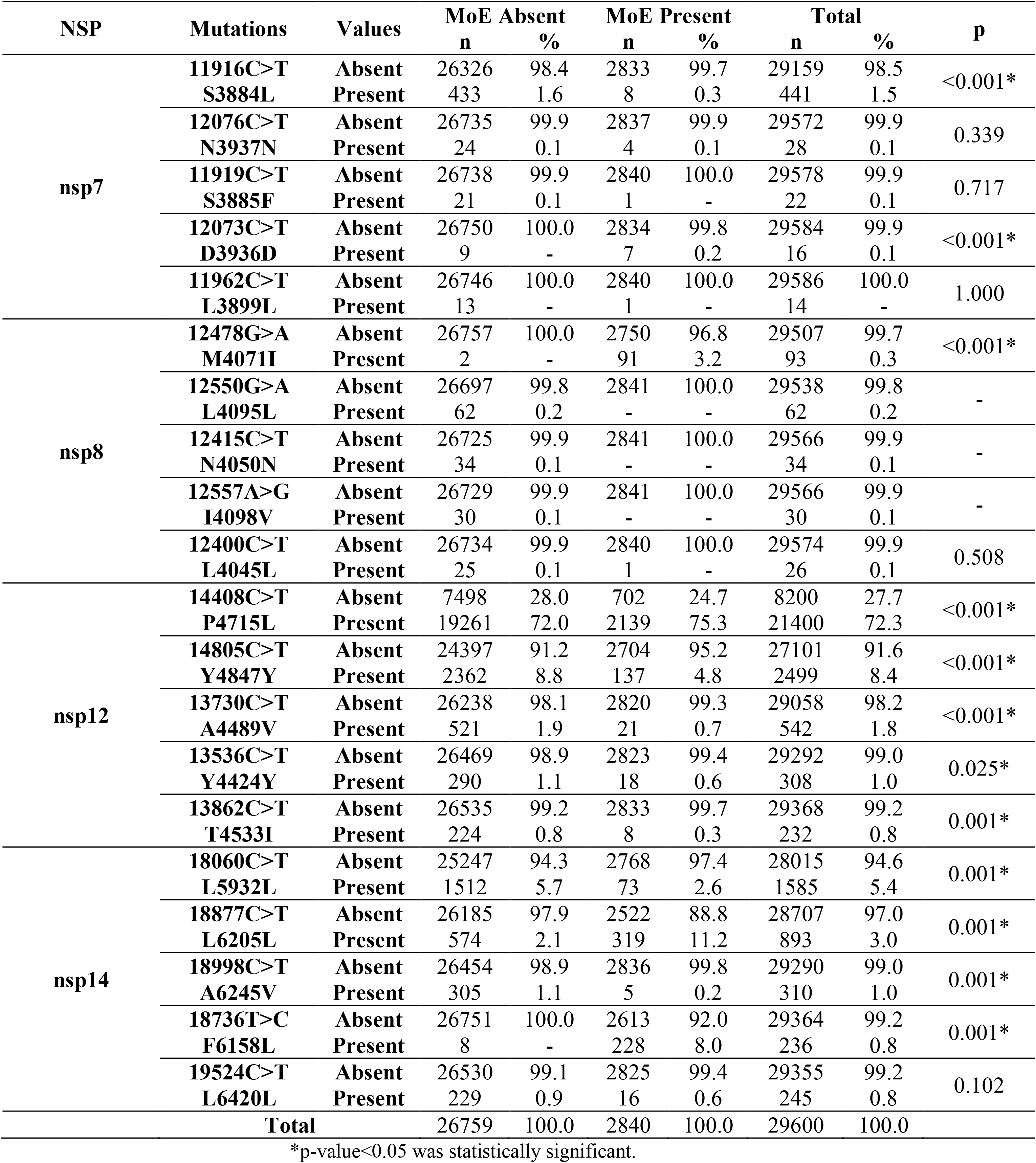
Comparisons of MoE and nsp mutations.

### Effects of geographical location on MoE status

In addition to time and genotype, we also examined the potential association between the location of isolates and MoE status as a possible confounding factor. We first examined whether there is a significant association between location, defined here as continent the isolate was originally obtained, and MoE status. Our results indicate that there is a strong association between location and MoE status, with the highest percentage of MoE present isolates in Asia (14.5%), and the percentage ratio in South America (6.5%) (p-value <0.001). In comparison to our previous findings, South America had a dramatic decrease in MoE present isolate percentage, likely as a result of the increased sequencing efforts (from 118 isolates to 416) removing potential sampling biases or localized founder effects. Africa, Asia, and North America had an increase in MoE present proportion, while Europe, Oceania, and South America showed lowered percentages (Table 2).

**Table 2.**
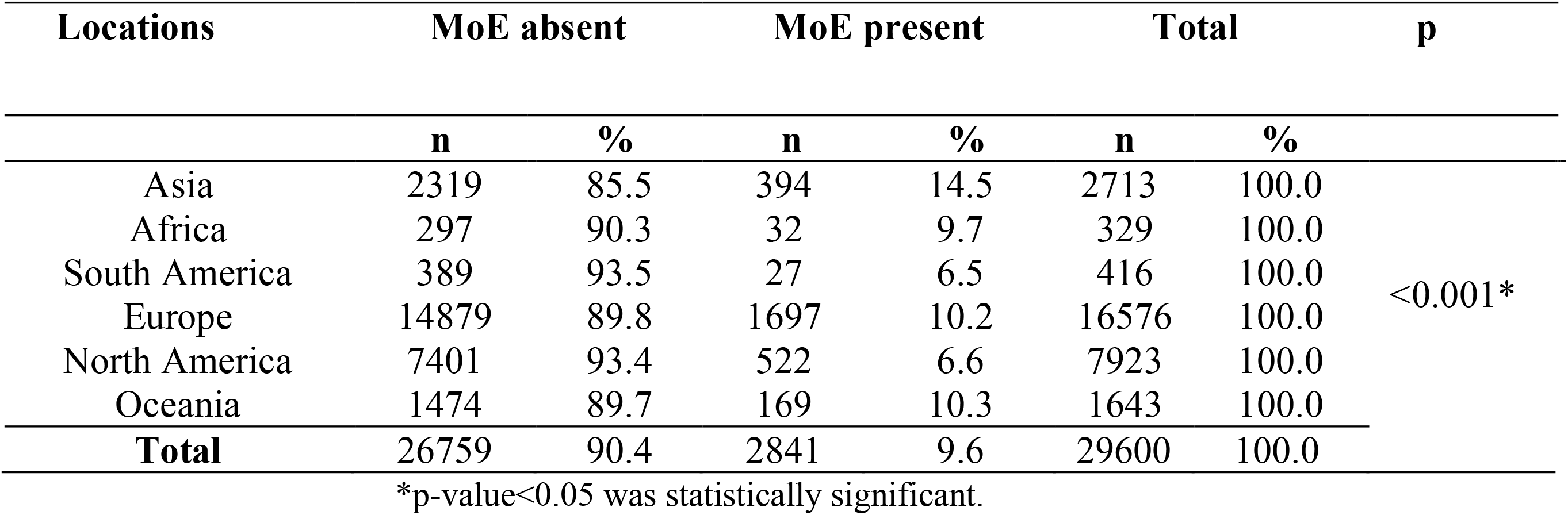
Distribution of MoE across geographical locations.

After observing the potential confounding effect of location on MoE status, we sought to understand whether a location is more or less likely to predict MoE status, using a logistic regression model (Table 3). Comparing each individual region (1) to the other five (0), we found that Asia, Europe, and North and South America are all possible predictors of MoE status (p-value < 0.05), with Asia and Europe 1.697 and 1.184 times as likely to be MoE present as the other regions, and North and South America 0.589 and 0.650 times as likely, respectively. Using these findings, we created different logistic regression models to identify which of the 12 mutations are likely to be independent predictors of MoE status (Table 4). In the single variable model, all 12 mutations we previously identified and location were found to be potential predictors (p-value < 0.05). Forming final models including the 12 mutations (Final Model A) and the mutations as well as locations (Final Model B), we observed that the predictor effect of two of the mutations nsp8 12478G>A and nsp14 18998C>T do not appear to be sufficiently independent of the other mutations in Final Model A. After adding the location variable to the Final Model A, location remains a significant predictor, with all five non-reference locations less likely to predict MoE than Asia, the reference location, and nsp12 14805C>T is found to not have a predictor effect independent of location (p-value = 0.073). Following Final Model B, nine mutations appear to have a significant association with MoE status, independent of other variables: 11916C>T, 12073C>T, 13536C>T, 13730C>T, 13862C>T, 14408C>T, 18060C>T, 18736T>C, and 18877C>T (p-value < 0.05).

**Table 3.** Logistic regression model of MoE and location on single variables. Each location was represented as itself (1) and others (0).

| Locations | p | OR | 95% C.I. |
| --- | --- | --- | --- |
| Asia | <0.001* | 1.697 | 1.513 to 1.903 |
| Africa | 0.937 | 1.015 | 0.703 to 1.465 |
| South America | 0.032* | 0.650 | 0.439 to 0.963 |
| Europe | <0.001* | 1.184 | 1.095 to 1.281 |
| North America | <0.001* | 0.589 | 0.533 to 0.650 |
| Oceania | 0.330 | 1.085 | 0.921 to 1.278 |

OR, Odds-Ratio; C.I.: confidence interval, \*p-value<0.05 was statistically significant.

**Table 4.**
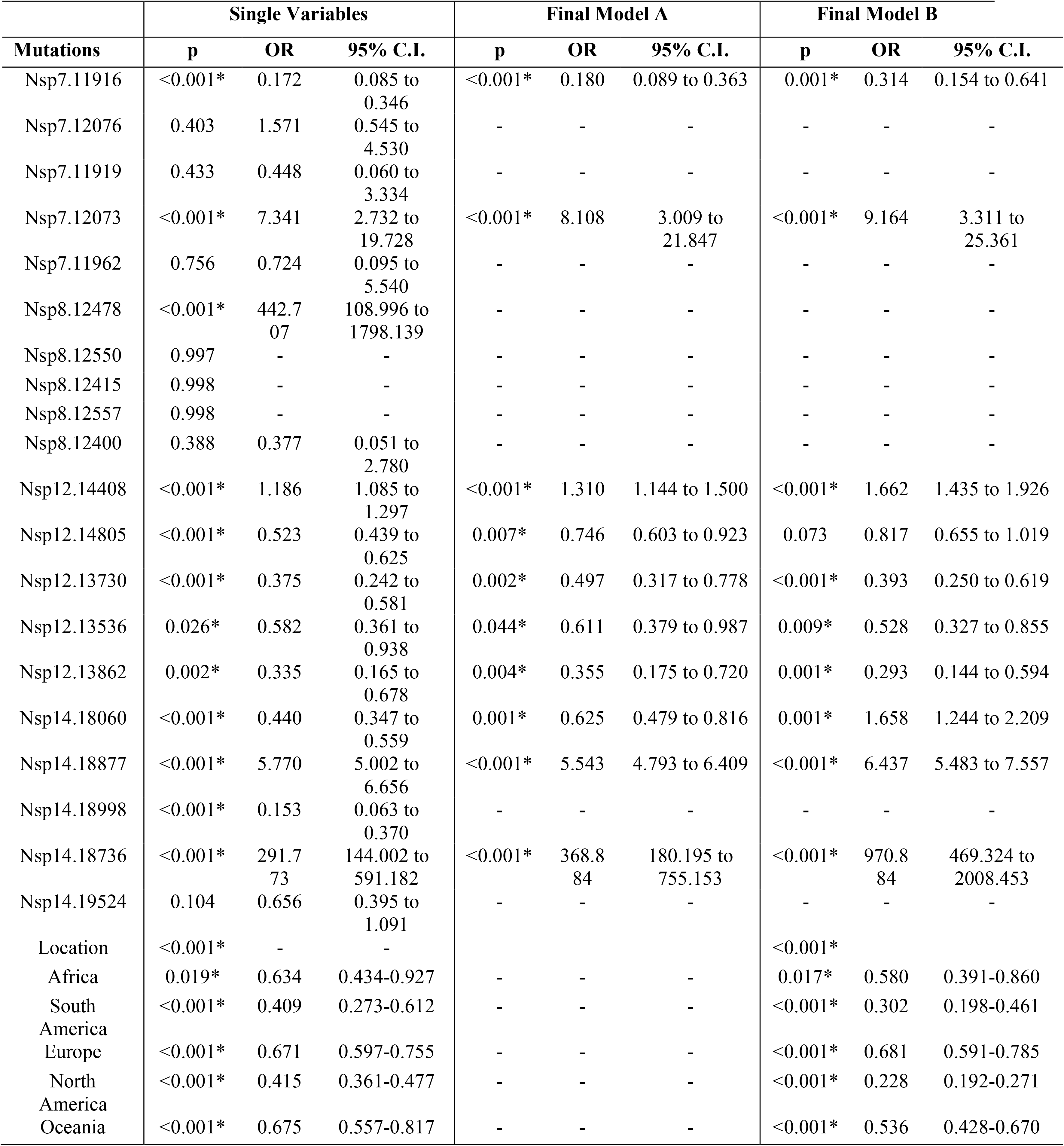
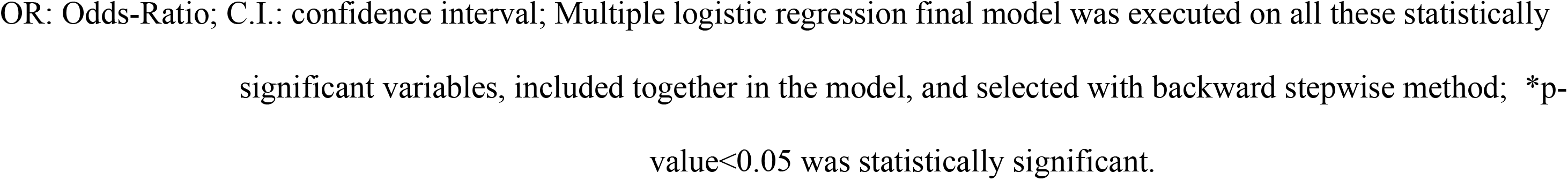
Logistic regression model of MoE on single variables and a final model. (Final Model A) Logistic regression model of ten mutations on final model. (Final Model B) Logistic regression model of four mutations and location on final model.

|  | Single Variables |  |  | Final Model A |  |  | Final Model B |  |  |
| --- | --- | --- | --- | --- | --- | --- | --- | --- | --- |
| Mutations | p | OR | 95% C.I. | p | OR | 95% C.I. | p | OR | 95% C.I. |
| Nsp7.11916 | <0.001* | 0.172 | 0.085 to 0.346 | <0.001* | 0.180 | 0.089 to 0.363 | 0.001* | 0.314 | 0.154 to 0.641 |
| Nsp7.12076 | 0.403 | 1.571 | 0.545 to 4.530 | - | - | - | - | - | - |
| Nsp7.11919 | 0.433 | 0.448 | 0.060 to 3.334 | - | - | - | - | - | - |
| Nsp7.12073 | <0.001* | 7.341 | 2.732 to 19.728 | <0.001* | 8.108 | 3.009 to 21.847 | <0.001* | 9.164 | 3.311 to 25.361 |
| Nsp7.11962 | 0.756 | 0.724 | 0.095 to 5.540 | - | - | - | - | - | - |
| Nsp8.12478 | <0.001* | 442.707 | 108.996 to 1798.139 | - | - | - | - | - | - |
| Nsp8.12550 | 0.997 | - | - | - | - | - | - | - | - |
| Nsp8.12415 | 0.998 | - | - | - | - | - | - | - | - |
| Nsp8.12557 | 0.998 | - | - | - | - | - | - | - | - |
| Nsp8.12400 | 0.388 | 0.377 | 0.051 to 2.780 | - | - | - | - | - | - |
| Nsp12.14408 | <0.001* | 1.186 | 1.085 to 1.297 | <0.001* | 1.310 | 1.144 to 1.500 | <0.001* | 1.662 | 1.435 to 1.926 |
| Nsp12.14805 | <0.001* | 0.523 | 0.439 to 0.625 | 0.007* | 0.746 | 0.603 to 0.923 | 0.073 | 0.817 | 0.655 to 1.019 |
| Nsp12.13730 | <0.001* | 0.375 | 0.242 to 0.581 | 0.002* | 0.497 | 0.317 to 0.778 | <0.001* | 0.393 | 0.250 to 0.619 |
| Nsp12.13536 | 0.026* | 0.582 | 0.361 to 0.938 | 0.044* | 0.611 | 0.379 to 0.987 | 0.009* | 0.528 | 0.327 to 0.855 |
| Nsp12.13862 | 0.002* | 0.335 | 0.165 to 0.678 | 0.004* | 0.355 | 0.175 to 0.720 | 0.001* | 0.293 | 0.144 to 0.594 |
| Nsp14.18060 | <0.001* | 0.440 | 0.347 to 0.559 | 0.001* | 0.625 | 0.479 to 0.816 | 0.001* | 1.658 | 1.244 to 2.209 |
| Nsp14.18877 | <0.001* | 5.770 | 5.002 to 6.656 | <0.001* | 5.543 | 4.793 to 6.409 | <0.001* | 6.437 | 5.483 to 7.557 |
| Nsp14.18998 | <0.001* | 0.153 | 0.063 to 0.370 | - | - | - | - | - | - |
| Nsp14.18736 | <0.001* | 291.773 | 144.002 to 591.182 | <0.001* | 368.884 | 180.195 to 755.153 | <0.001* | 970.884 | 469.324 to 2008.453 |
| Nsp14.19524 | 0.104 | 0.656 | 0.395 to 1.091 | - | - | - | - | - | - |
| Location | <0.001* | - | - |  |  |  | <0.001* |  |  |
| Africa | 0.019* | 0.634 | 0.434-0.927 | - | - | - | 0.017* | 0.580 | 0.391-0.860 |
| South America | <0.001* | 0.409 | 0.273-0.612 | - | - | - | <0.001* | 0.302 | 0.198-0.461 |
| Europe | <0.001* | 0.671 | 0.597-0.755 | - | - | - | <0.001* | 0.681 | 0.591-0.785 |
| North America | <0.001* | 0.415 | 0.361-0.477 | - | - | - | <0.001* | 0.228 | 0.192-0.271 |
| Oceania | <0.001* | 0.675 | 0.557-0.817 | - | - | - | <0.001* | 0.536 | 0.428-0.670 |

OR: Odds-Ratio; C.I.: confidence interval; Multiple logistic regression final model was executed on all these statistically significant variables, included together in the model, and selected with backward stepwise method; \*p-value<0.05 was statistically significant.

### Nsp14 mutations have significant impact on increased genomic mutation density

We then examined the effects of each mutation on genomic mutation density to see whether the relationship between the mutations and MoE status are indicative of a genome-wide trend. Due to selection potentially effecting nonsynonymous mutations differentially, we separated the mutations in the two categories and calculated mutation density separately for each category. Our results show that nsp14 mutations show the most consistent association with mutations between MoE and the whole genome. All three nsp14 mutations (18060C>T, 18736T>C, and 18877C>T) which have a significant association with MoE status also show a similar relationship with genomic mutation density (Fig. 2). 18060C>T (L7L) has the lowest odds ratio for MoE status (Table 4), and while it shows a slower increase in synonymous mutation density compared to wildtype isolates (Fig. 2A), it has a significant impact on faster mutation density increase in nonsynonymous mutations (Fig. 2B). In comparison, 18877C>T (L270L) (Fig. 2C-D) and 18736T>C (F233L) (Fig. 2E-F) both show a high prediction capacity for MoE and an increased mutation density. In comparison, mutations in nsp7 (Supp. Figs. 1-2) and nsp12 (Supp. Figs. 3-6) show much lower impact on altered mutation density increase rate. 12073C>T, an nsp7 mutation, displays high divergence from wildtype isolate patterns; however, its low sample size (n = 16) creates a skewed distribution of isolates across time, complicating any potential inference. It bears noting that not all mutations continue to be observed up to the date of retrieval. 18060C>T and 18736T>C have been last observed on 7 May and 11 May, 2020, respectively. Whether this is due to sequence sampling or because the mutations confer an evolutionary disadvantage is unknown. 18060C>T and 18877C>T are both synonymous mutations, yet only the latter continues to be present in recent isolates.

**Figure 2.**
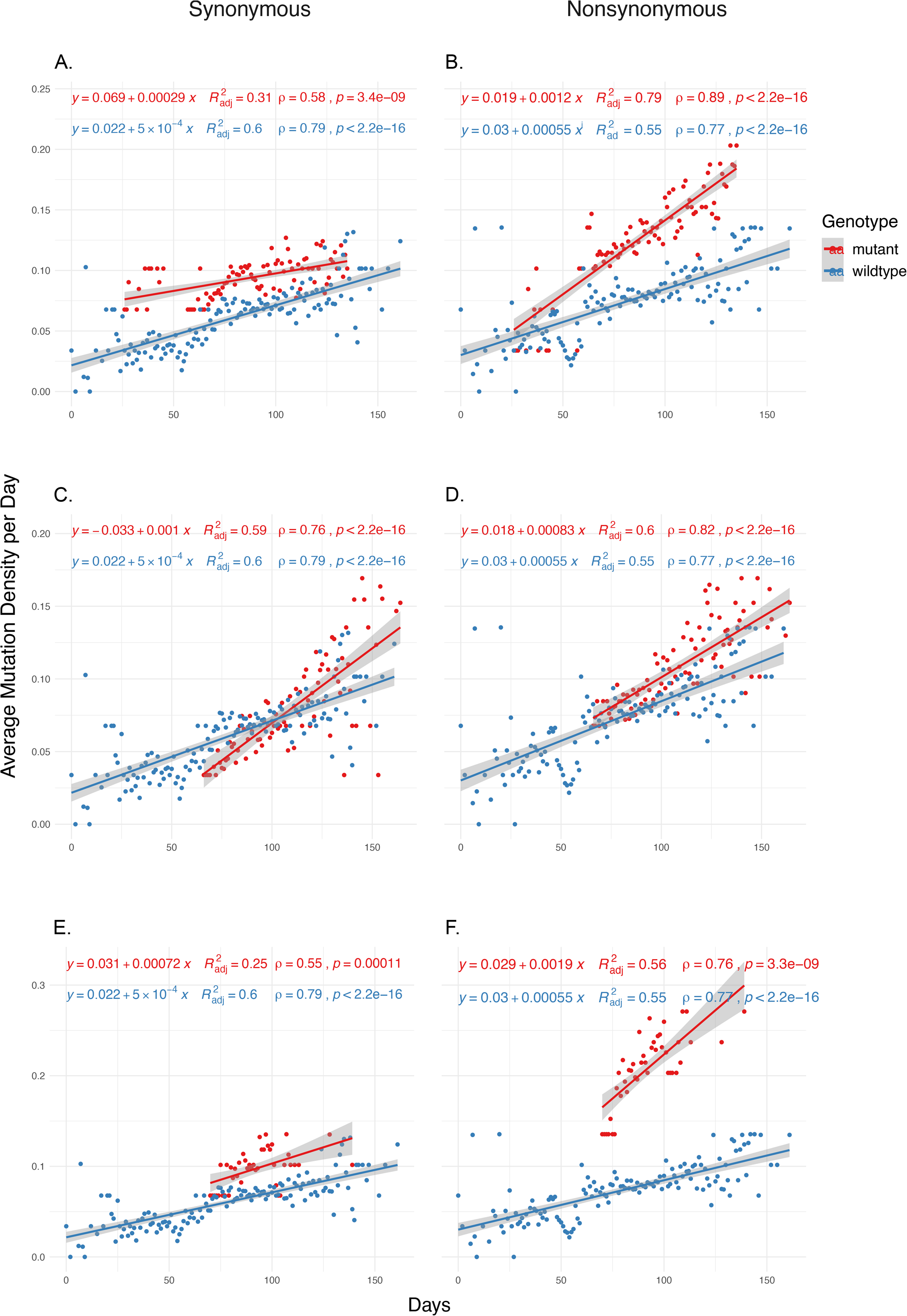
The distribution of synonymous and nonsynonymous mutations in isolates carrying nsp14 mutations compared to wildtype isolates. (A-B) Isolates carrying the synonymous18060C>T mutation (n = 1585). (C-D) Isolates carrying the synonymous 18877C>T mutation (n = 893). (E-F) Isolates carrying the nonsynonymous 18736C>T mutation (n=236). Wildtype isolates in all graphs carry the reference nucleotide for the nine positions of interest (11916, 12073, 13536, 13730, 13862, 14408, 18060, 18736, 18877) (n = 5910). Correlation scores are calculated using Spearman rank correlation.

## Conclusions

Our previous work identified RdRp mutations as contributors to the evolution of the SARS-CoV-2 genome and this study confirmed those findings. Furthermore, we hypothesized that mutations of the other critical components of the viral replication and transcription machinery may have similar effects. Our results implicate nsp14 as a source of increased mutation rate in SARS-CoV-2 genomes. Three of the five most common nsp14 mutations, namely 18060C>T, 18736C>T and 18877C>T are associated with increases in both genome-wide mutational load, as well as MoE status, an alternative indicator of mutational rate and virus evolution. Interestingly all three are located within the ExoN domain, which is responsible for the proofreading activity of nsp14; however, only 18736C>T mutation is non-synonymous (F233L), while 18060C>T and 18877C>T are synonymous mutations and therefore, only after functional studies it will be possible to understand their effects on viral replication processes.

The fate of three nsp14 mutations are also intriguing: Despite being present in the first case detected in the Washington state of the US in mid-January, and detected in 1,585 cases till May 7, 18060C>T mutation has not been detected since then. On the other hand, 18877C>T mutation arising around at the end of January likely in Saudi Arabia and being detected in much less cases (n=893), is still present in many isolates. However, it should be noted that 18877C>T mutation arose within the dominant 23403A>G / 14408C>T lineage, while the other two nsp14 mutations are in different lineages. Therefore, dominance or disappearance of different nsp14 mutations may have less to do with these particular mutations and more with the co-mutations. Yet, we cannot rule out possible effects of these nsp14 mutations on the fitness of SARS-CoV-2.

## Additional Information and Declarations

## Acknowledgement

The authors would like to thank Mr. Alirıza Arıbaş from Izmir Biomedicine and Genome Center for his technical assistance. The authors also would like to extend their thanks to Izmir Biomedicine and Genome Center (IBG) COVID19 platform IBG-COVID19 for their support in implementing the study and the Scientific and Technological Research Council of Turkey (TUBITAK) for their financial support of IBG-COVID19.

## Grant Disclosures

The following grant information was disclosed by the authors:

Turkish Academy of Sciences Young Investigator Program (TÜBA-GEBİP).

## Author Contributions

Doğa Eskier, Aslı Suner, Gökhan Karakülah and Yavuz Oktay conceived and designed the experiments, performed the experiments, analyzed the data, prepared figures and/or tables, authored or reviewed drafts of the paper, and approved the final draft.

## Data Availability

The data is available at Mendeley: Eskier, Doğa; Suner, Aslı; Oktay, Yavuz; Karakülah, Gökhan (2020), “SARS-CoV-2 GISAID isolates (2020-06-17) genotyping VCF”, Mendeley Data, v1. http://dx.doi.org/10.17632/63t5c7xb4c.1

## Supplemental Information

Supplemental materials are included with this research.

