## Supplemental Figures for "Mutations of SARS-CoV-2 nsp14 exhibit strong association with increased genome-wide mutation load"

#### Supplementary Figure 1

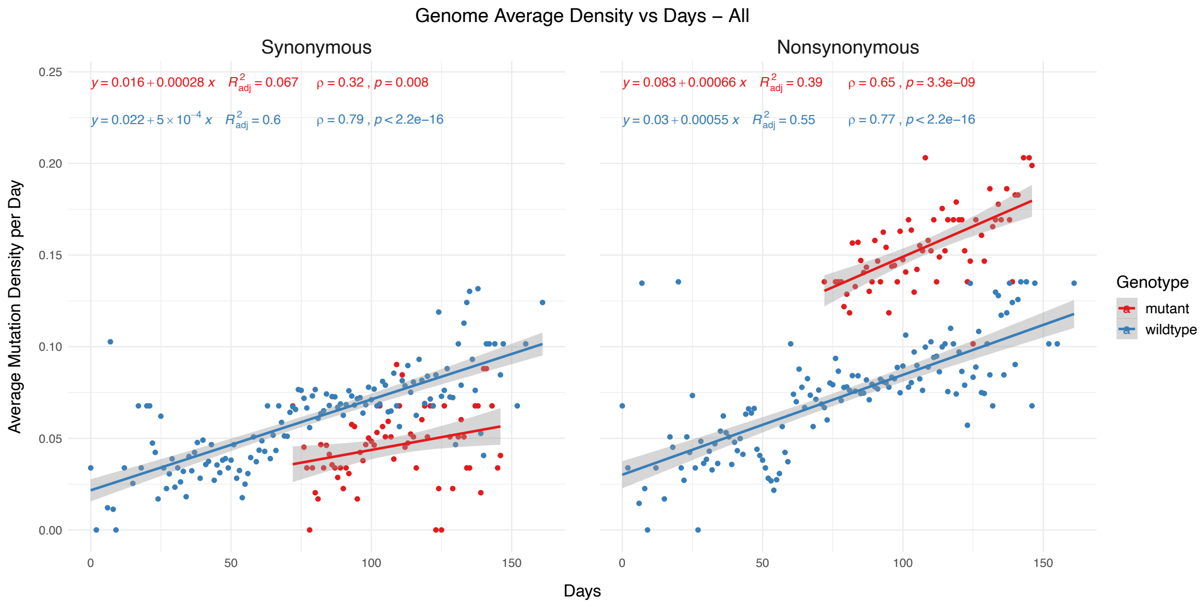


**Supplementary Figure 1. The distribution of synonymous and nonsynonymous mutations in isolates carrying 11916C>T mutation compared to wildtype isolates**.

Correlation scores are calculated using Spearman rank correlation.

#### Supplementary Figure 2

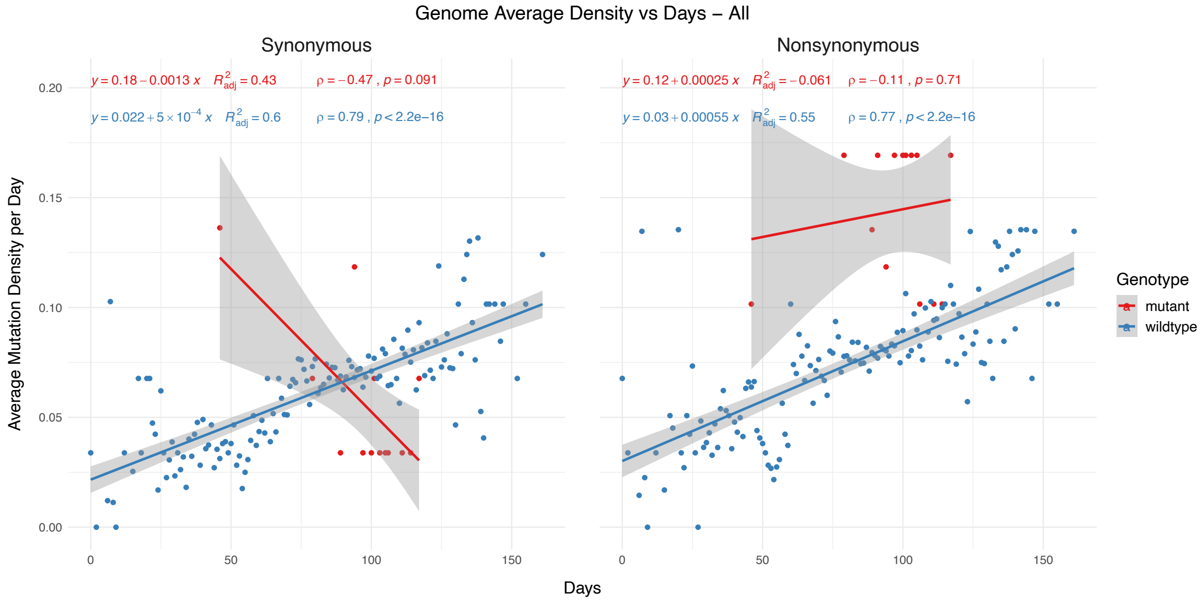


**Supplementary Figure 2. The distribution of synonymous and nonsynonymous mutations in isolates carrying 12073C>T mutation compared to wildtype isolates**.

Correlation scores are calculated using Spearman rank correlation.

### Supplementary Figure 3

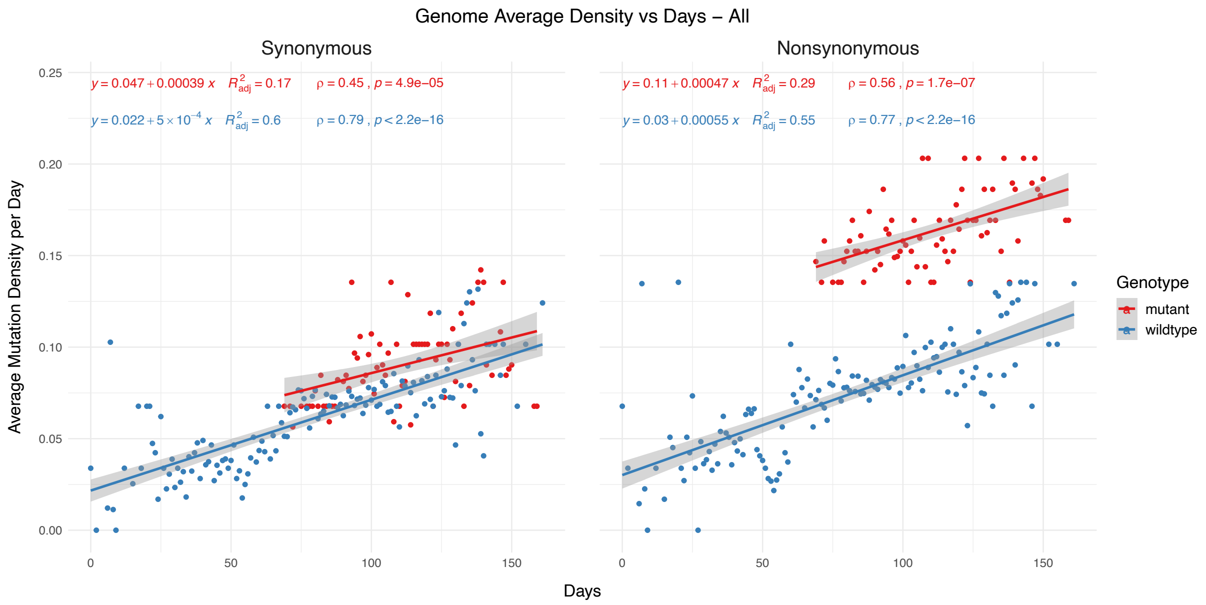


**Supplementary Figure 3. The distribution of synonymous and nonsynonymous mutations in isolates carrying 13536C>T mutation compared to wildtype isolates**.

Correlation scores are calculated using Spearman rank correlation.

#### Supplementary Figure 4

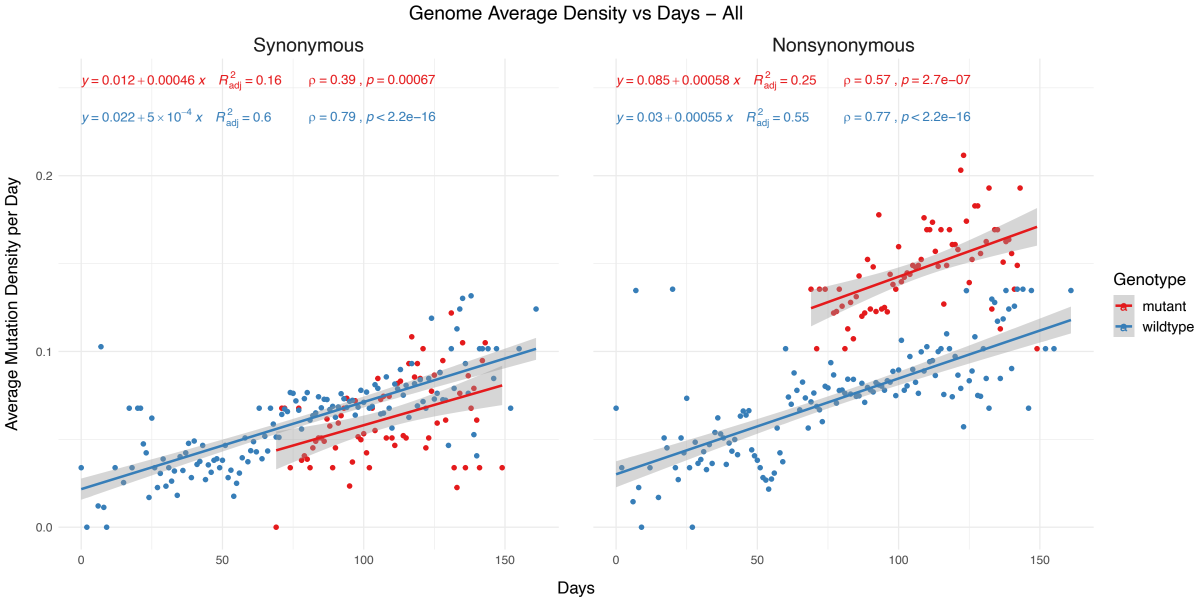


**Supplementary Figure 4. The distribution of synonymous and nonsynonymous mutations in isolates carrying 13730C>T mutation compared to wildtype isolates**. Correlation scores are calculated using Spearman rank correlation.

### Supplementary Figure 5

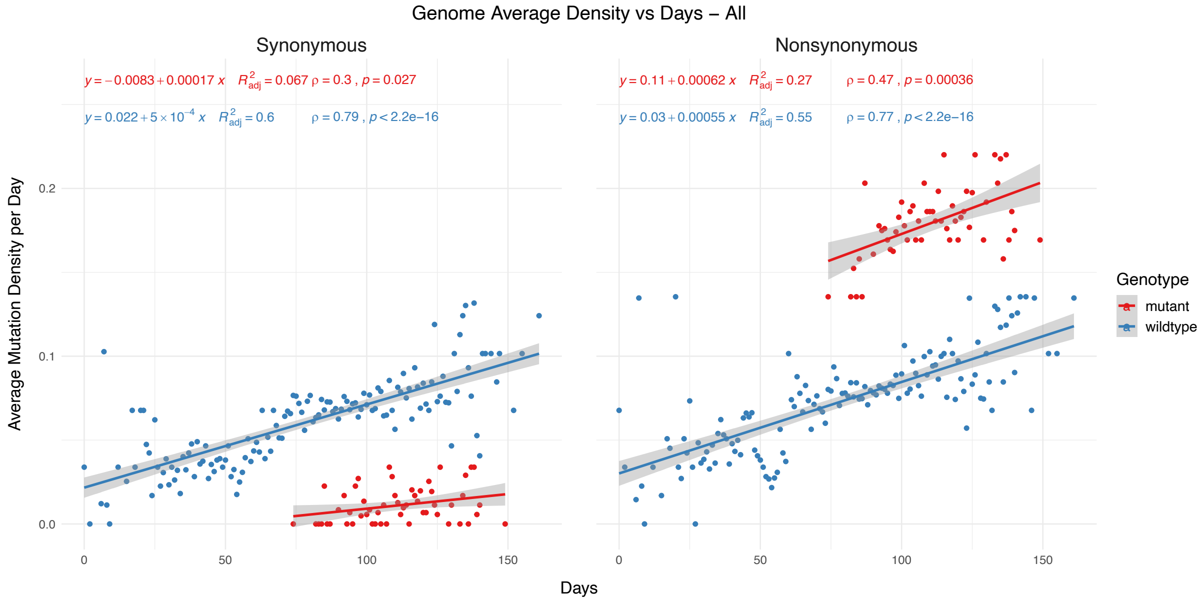


**Supplementary Figure 5. The distribution of synonymous and nonsynonymous mutations in isolates carrying 13862C>T mutation compared to wildtype isolates**.

Correlation scores are calculated using Spearman rank correlation.

#### Supplementary Figure 6

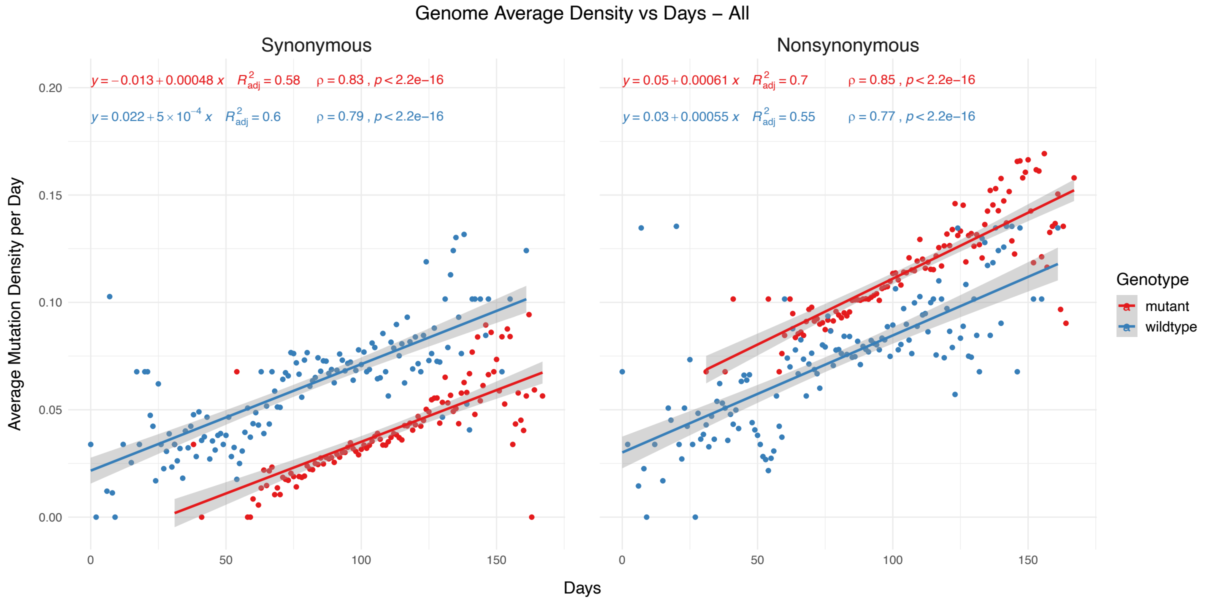


**Supplementary Figure 6. The distribution of synonymous and nonsynonymous mutations in isolates carrying 14408C>T mutation compared to wildtype isolates**.

Correlation scores are calculated using Spearman rank correlation.
